# Photoemission electron microscopy for connectomics

**DOI:** 10.1101/2023.09.05.556423

**Authors:** Kevin M. Boergens, Gregg Wildenberg, Ruiyu Li, Lola Lambert, Amin Moradi, Guido Stam, Rudolf Tromp, Sense Jan van der Molen, Sarah B. King, Narayanan Kasthuri

**Affiliations:** Department of Physics, University of Illinois Chicago; Department of Neurobiology, University of Chicago; Argonne National Laboratory; Department of Chemistry, University of Chicago; James Franck Institute, University of Chicago; Leiden Institute of Physics, Leiden University; IBM T.J. Watson Research Center

## Abstract

Detailing the physical basis of neural circuits with large-volume serial electron microscopy (EM), ‘connectomics’, has emerged as an invaluable tool in the neuroscience armamentarium. However, imaging synaptic resolution connectomes is currently limited to either transmission electron microscopy (TEM) or scanning electron microscopy (SEM). Here, we describe a third way, using photoemission electron microscopy (PEEM) which illuminates ultra-thin brain slices collected on solid substrates with UV light and images the photoelectron emission pattern with a wide-field electron microscope. PEEM works with existing sample preparations for EM and routinely provides sufficient resolution and contrast to reveal myelinated axons, somata, dendrites, and sub-cellular organelles. Under optimized conditions, PEEM provides synaptic resolution; and simulation and experiments show that PEEM can be transformatively fast, at Gigahertz pixel rates. We conclude that PEEM imaging leverages attractive aspects of SEM and TEM, namely reliable sample collection on robust substrates combined with fast wide-field imaging, and could enable faster data acquisition for next-generation circuit mapping.

## 2 Introduction

A confluence of advances in 3D EM imaging, sample preparation, and algorithms has enabled the exhaustive mapping of how neurons connect in large volumes of brain – connectomics. Connectomics has emerged as an invaluable tool in the neuroscience toolbox and the analysis of wiring patterns in cortex, hippocampus, retina, songbird sensory motor cortex, zebrafish larvae, and entire fly brains have helped discover principles of brain functions that could not have been revealed in any other way [1, 2, 3, 4, 5, 6, 7, 8]. These datasets are also inherently valuable as public repositories for future investigations by the broader neuroscience community [9, 10]. Thus, there is only increased demand for even larger 3D EM datasets. The next leap in connectomics will be reconstructing full neural circuits in mammalian brains. But to make that leap, large-volume connectomes need to be cheaper and faster. A 50-100 fold drop in the price of a ‘connectomic voxel’ could revolutionize connectomics, similar to the way a 50-100 fold reduction in the cost of DNA sequencing [11] placed large-scale DNA analysis at the core of modern biology. However, all EM connectomic reconstructions use either transmission electron microscopy (TEM) or scanning electron microscopy (SEM) [12], which have limitations in the reliable automated serial pickup of ultrathin brain slices (UTBS), or in acquisition costs, respectively.

In this study we investigated whether photoemission electron microscopy (PEEM) would be a third option. Photoemission electron microscopes are commercially available, are routinely used in chemistry and surface physics [13, 14, 15, 16, 17] and have been used since the 1970s for biological imaging [18, 19, 20], and even to image neurons in culture [21]. PEEM uses wide-field illumination (such as UV light) to emit photoelectrons from materials and captures variations in the resulting photoemission pattern with standard electron optics [22] (**Figure 1**). Photoemitted electrons are accelerated (usually to 10 keV to 20 keV) and projected with electrostatic and electromagnetic lenses onto an electron detector. Commercial PEEMs can have resolution capabilities of less than 15 nm, which is suitable for synaptic connectomics [23]. There are also cameras available that capture PEEM data at multi-GHz pixel rates, making a fast PEEM for connectomics plausible. Here, we report the first study of PEEM imaging of ultrathin brain slices prepared using standard connectomics sample preparation methods[24].

**Figure 1:**
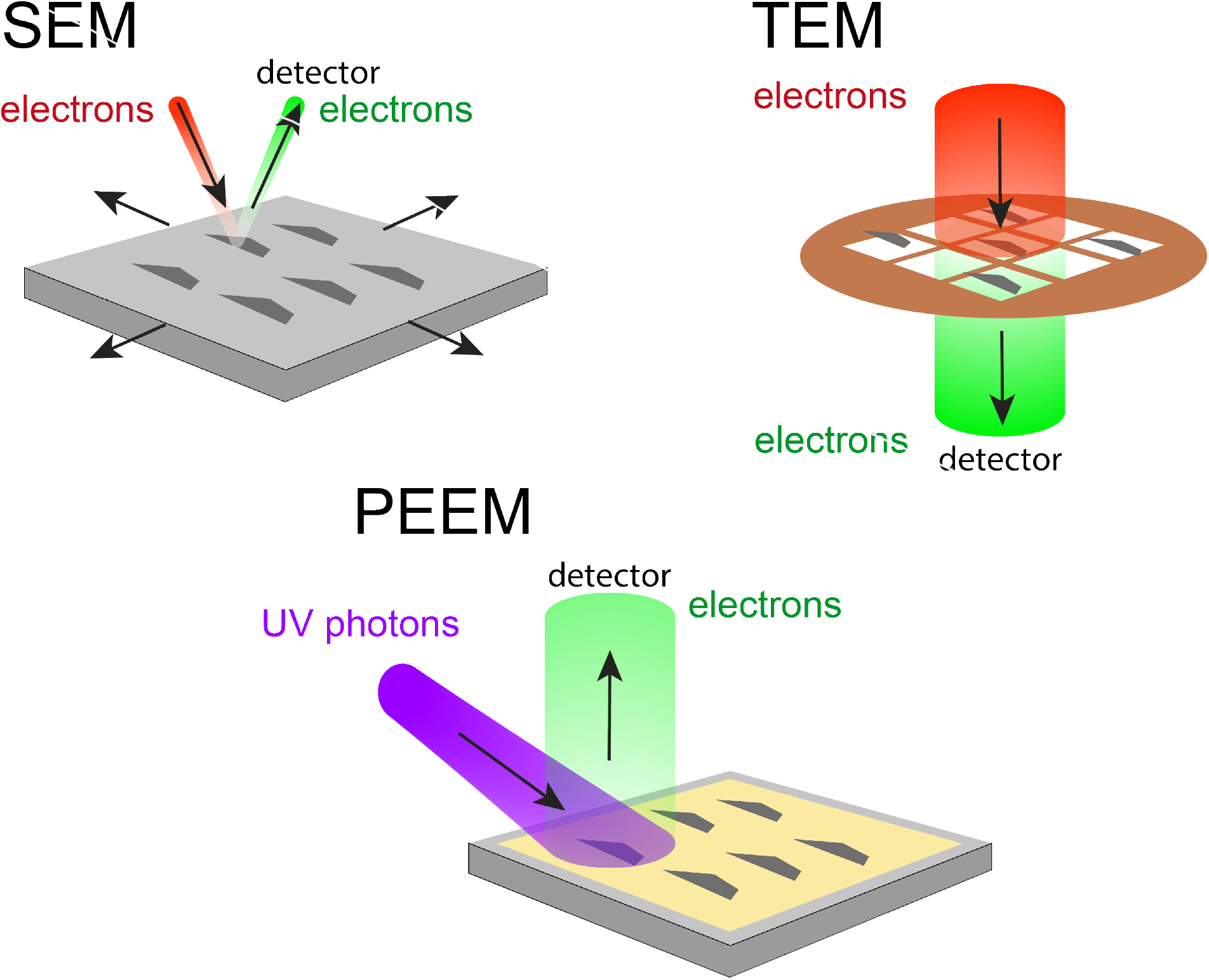
Photoemission electron microscopy (PEEM) combines the advantages of SEM and TEM for high-throughput volume electron microscopy (vEM). SEM allows for more reliable section collection on solid substrates, but is slower due to the inherently sequential nature of scanning microscopy. TEM is wide-field imaging and fast but samples are collected onto grids with thin fragile substrates, increasing the complexity of collecting thousands of UTBS. PEEM combines the best of both, imaging samples on solid substrates with wide-field imaging technique, and thus has the potential to supersede these techniques and enable the next generation of large-volume connectomes and projectomes.

## 3 Results

As PEEM is usually done on flat conductive substrates, we manually picked up 60 nm brain sections that had been prepared for EM-based connectomics [24] on a square-centimeter-sized piece of silicon, coated with a 50 nm layer of gold, and glow-discharged to increase hydrophilicity and adhesiveness of brain sections. We found that UTBS could be reliably imaged in a PEEM with a mercury arc lamp. We were able to identify biological features readily observed in standard connectomic datasets including cellular features (e.g., somata, blood vessels, dendrites, and myelinated axons) and sub-cellular features (e.g., mitochondria, endoplasmic reticulum, nuclear membrane, and nucleolus) (**Figure 2**). PEEM imaging is also relatively free of distortions – we tiled multiple overlapping PEEM fields of view (FOV) and collected and aligned small stacks of PEEM images of serial sections (**Figure 3**). In such stacks, we traced the processes of myelinated axons, dendrites, and glia, demonstrating that PEEM works in 3D for volume electron microscopy (vEM) and can be used to reconstruct projectomes [25]. Lastly, we acquired a high-resolution resolution PEEM image where synapses could be clearly identified with individual, putative *∼* 40 nm diameter, synaptic vesicles (**Figure 4**), meaning PEEMs can deliver sufficient resolution for full connectomic circuit reconstruction.

**Figure 2:**
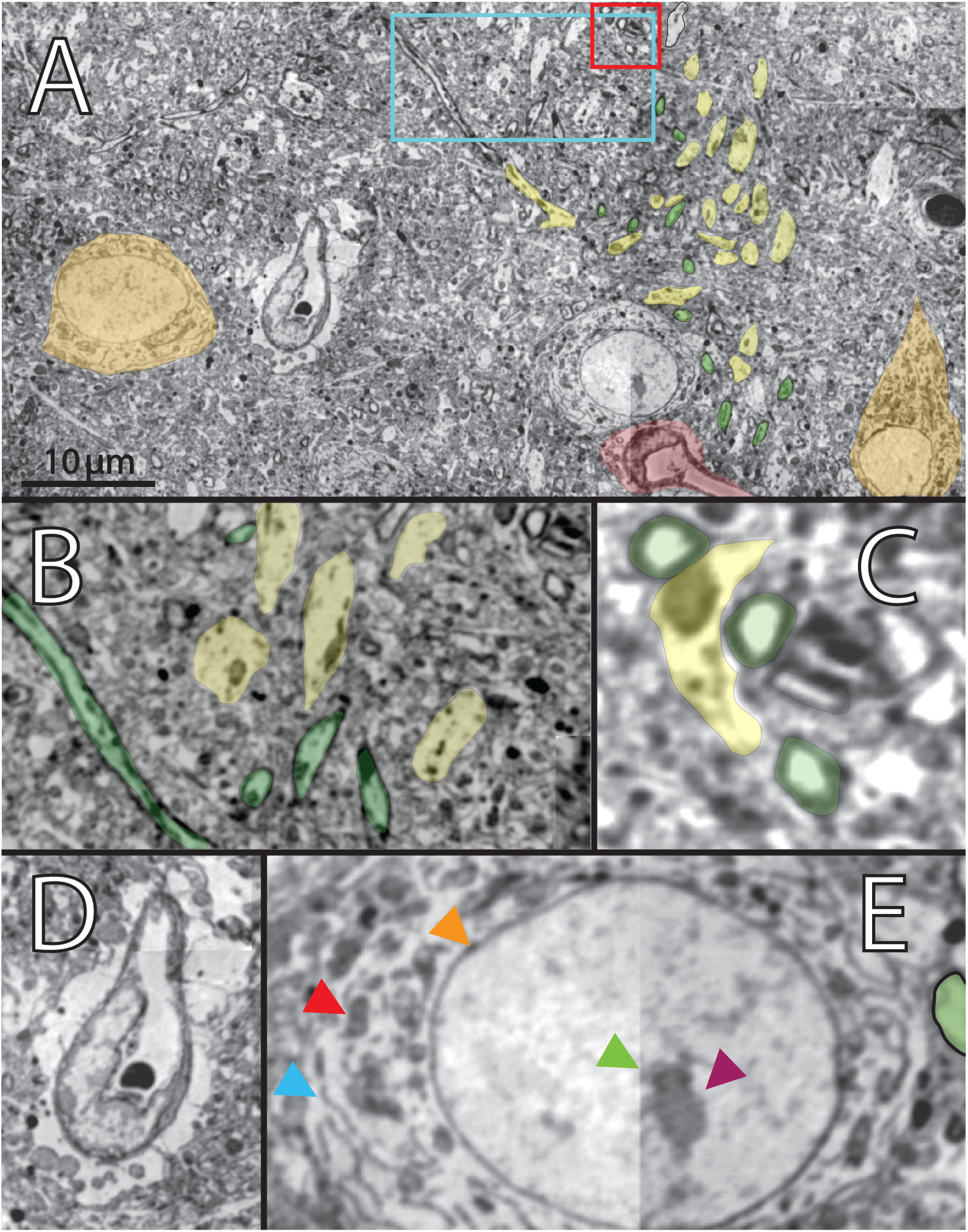
PEEM reveals cellular and subcellular features similar to standard EM data. **A)** a 60 nm UTBS was imaged over 9 overlapping fields of view (FOV) spanning 72 μm × 37 μm and algorithmically stitched together. PEEM images clearly show different neurobiological features commonly found in EM datasets including somata (orange), blood vessels (red), dendrites (yellow), and individual myelinated axons (green). **B)** Zoom in from A (light blue box) showing close-up of individual dendrites (yellow) and myelinated axons (green). **C)** Zoom in from B (red box) showing further detail of individual dendrites (yellow) and myelinated axons (green). **D)** Representative example of a blood vessel with the processes of an astrocyte making a putative neurovascular unit. **E)** Representative example of a neuronal soma with clearly visible mitochondria (red arrowhead), endoplasmic reticulum (light blue arrowhead), nucleolus (purple arrowhead), and nuclear membrane (orange arrowhead). Green arrowhead shows stitch line between two FOVs.

**Figure 3:**
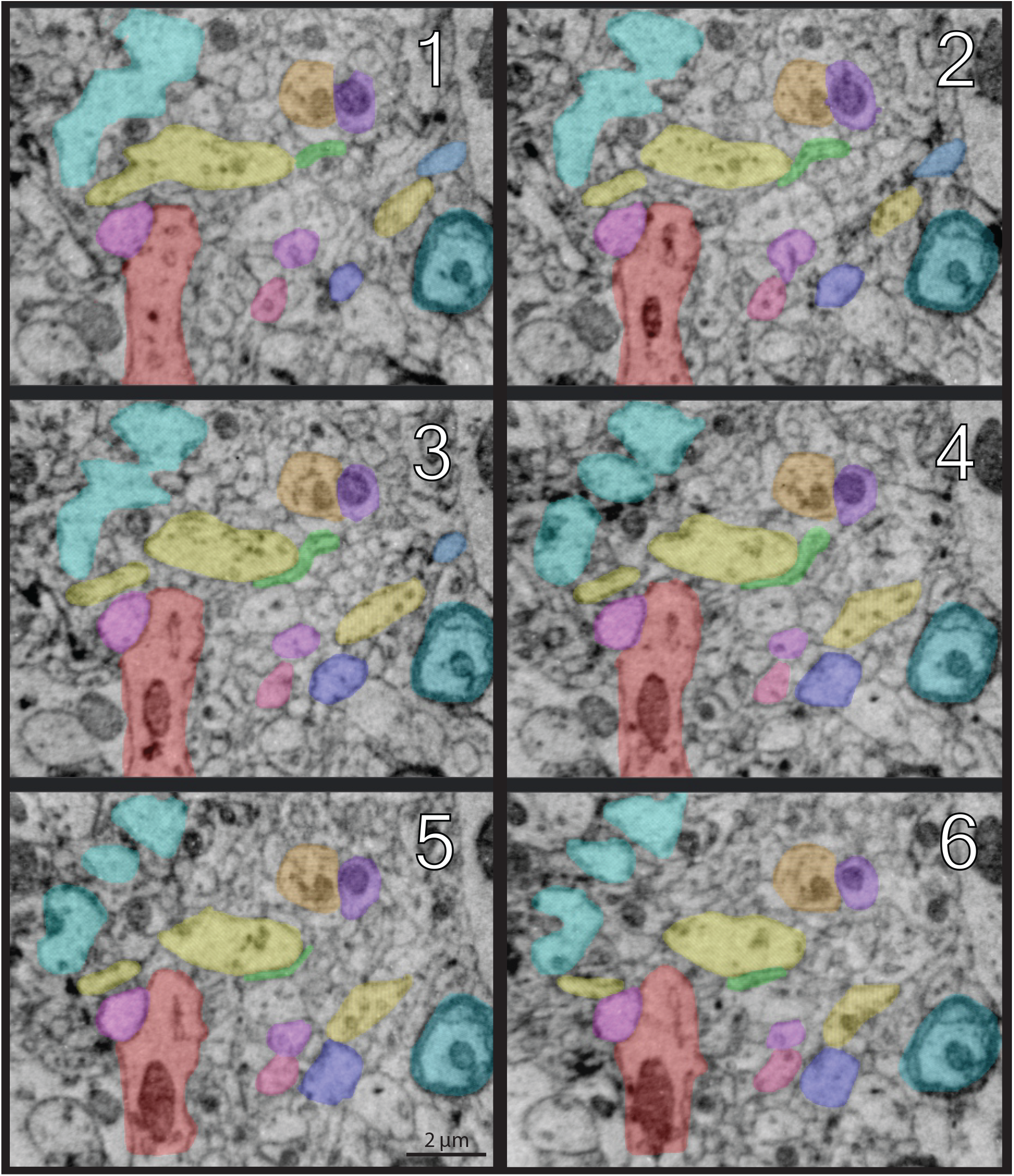
Tracing neurites in a serial stack of UTBS imaged with PEEM. Shown are images of a serial stack of UTBS, 40 nm thick, imaged with medium-resolution PEEM, suitable for fast projectomics. The stack was aligned using AlignTK. Multiple neurological features can be manually traced through the stack including dendrites of varying size, myelinated axons, and glia across 6 sections.

**Figure 4:**
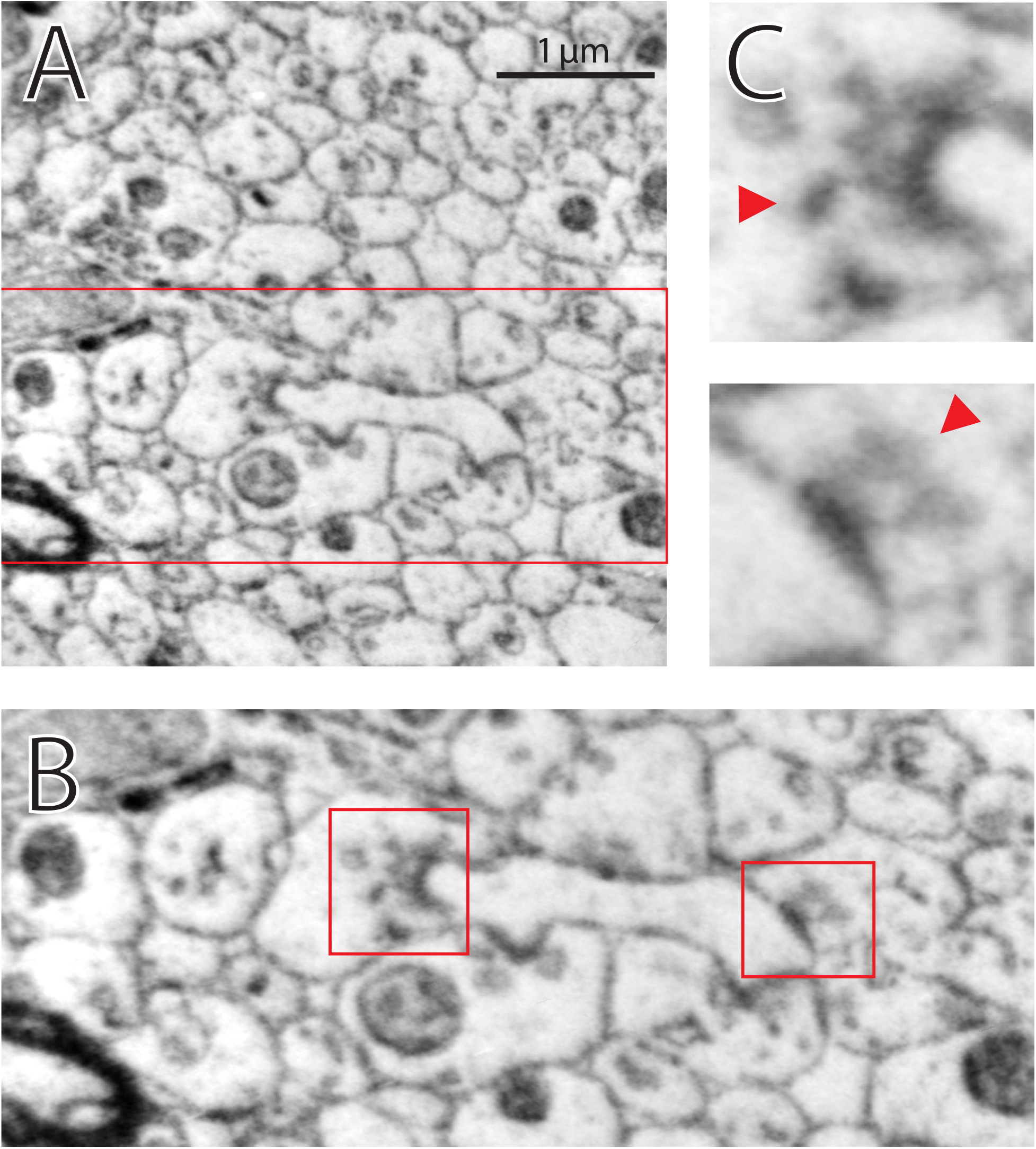
PEEM can achieve synaptic resolution. A) High-resolution PEEM image of a UTBS. Zoom in from A (red rectangle) showing multiple synapses (red squares) as evidenced by the presence of a post-synaptic densities (PSD) and pre-synaptic vesicle clouds. C) Zoom in from B (red squares) of two synapses with individual pre-synaptic vesicles visible (red arrowheads).

We next explored whether PEEM imaging could be fast, potentially at Gigahertz acquisition rates. For the results shown in this study, mercury arc lamps were used to illuminate the sample but the low photon flux necessitates long integration times (i.e., minutes) which is too slow to image large volumes with PEEM. An obvious upgrade would be to illuminate instead with UV lasers which trivially produce an average photon flux over 10,000 times higher than arc lamps. Thus, we investigated the compatibility of UTBS with laser-based illumination.

We first conservatively estimated that at most 3 million photons are needed to elicit an electron (see **Methods**). We then measured the emissivity ratio between the sample regions with the lightest and heaviest osmium stained structures, i.e. the cytoplasm versus a mitochondria in **Figure** 2, and found a ratio of 2.7 *±* 0.2. A 0.5 W 244 nm laser produces a flux of 6.1 × 10^17^ photons per second and using the Rose contrast criterion [26], we concluded that such a laser would provide enough signal to support connectomic resolution imaging at a pixel rate of over 2 GHz. Notably, CW lasers of such specifications are commercially available.

We next asked whether such illumination, i.e. a 0.5 W laser focused onto a field of view (FOV) tens of microns wide, would damage UTBS. We measured the absorption of UTBS for a range of wavelengths (**Supplementary Figure 1**). Interestingly, the absorption in epoxy has a local minimum around 250 nm, suggesting that wavelengths lower than that may be more efficient for imaging (see Discussion). Using this absorption data, we simulated the heating of a 40 nm thick brain slice on a reflective silicon substrate with a 40 μm × 40 μm FOV by a 0.5 W laser and found that this would heat the epoxy by only 21.6 K (**Supplementary Table 1**). We experimentally confirmed (using a 266 nm Raman CW laser) that a slice of epoxy on silicon can withstand the equivalent photon flux (0.31 mW/μm^2^) without epoxy burn-off. In anticipation of future development of more powerful UV-C lasers, we also simulated UTBS on more thermally conductive substrates, copper and diamond, and found that they would support 1.6 W and 3.9 W lasers respectively without going beyond the UTBS heating described above. This would allow PEEMs to image at 6.4 and 15.6 Gigahertz pixel rate, respectively. Lastly, we simulated space charging, i.e., whether electrons emitted at Gigahertz rates from 10’s of microns FOVs would potentially interact with each other, repulsing each other and reducing the effective resolution of PEEM imaging, which has been cited as a potential limit of PEEM imaging speed [27, 28]. We found that PEEM imaging with a 0.5 W CW laser and a 40 μm × 40 μm FOV deflected electrons no more than 1 nm. At 15.6 GHz pixel rate, 0.2 % of electrons would deflect more than that (**Supplementary Figure 2**, a result in good agreement with prior work on space charging in PEEM [29]). Only at sample current densities equivalent to a throughput of hundreds of Gigahertz space charging becomes a concern. This buffer means that non-CW lasers would also be compatible with fast, well-resolved connectomic data acquisition, as long as for a majority of the time electrons are in-flight. We conclude that Photoemission electron microscopy holds significant promise for high-throughput brain mapping and adjacent fields.

## 4 Discussion

Here we describe the first successful imaging of UTBS using PEEM at synaptic resolution and routine image acquisition at projectomic resolution. Furthermore, we demonstrate the plausibility of using UV laser illumination of UTBS to significantly increase PEEM imaging rates, exceeding state-of-the-art SEM and TEM techniques used for connectomic imaging of brains. Below we discuss future directions on how to make large-volume connectomic and projectomic PEEM imaging possible, with the acquisition of the data for a whole mouse brain within reach.

### 4.1 A PEEM mouse brain

An adult mouse brain, conservatively estimated to be 1 cm^3^, imaged at 15 nm isotropic resolution, as tentatively demonstrated above, would result in a dataset of 2.96 × 10^17^ voxels. A single PEEM with a CW UV laser, imaging non-stop at 2 Gigahertz pixel rate, would take approximately five years to finish that acquisition. UTBS can be easily distributed across multiple PEEMs, decreasing the time to map a mouse brain proportionately to the number of PEEMs used. A 2 GHz PEEM is feasible given the analyses above and would be faster than current approaches using TEMs or SEMs with a potentially significant reduction in price.

Another application of PEEMs would be to map, at lower resolution, projectomes of entire mouse brains, with imaging times of a few months. We found imaging at 40 nm projectomic resolution to work reliably; and such mapping makes good use of the large field of view of PEEM setups, which can be up to 800 μm. Finally, we found preliminary evidence that SOA machine learning algorithms for identifying objects in scenes could be easily trained to identify cell boundaries in PEEM datasets (Segment Anything from Meta [30], Supplementary Figure 3). While not surprising, given the similarity of PEEM brain images to SEM and TEM, such preliminary data suggests that a PEEM mouse projectome can be both quickly acquired and reconstructed in the near future.

PEEM connectomes and projectomes will require automating both the collection and imaging of 1000s of UTBS. Since PEEM has similar requirements as SEM imaging (i.e., section flatness, conductive substrates, and maximizing UTBS density), we argue existing state-of-the-art approaches for automated collection of serial UTBS on conductive substrates, such as automated collection on tape (ATUM) [4, 31, 32] and collection on silicon, MagC [33, 34] could be used to scale the number of samples for PEEM.

We end with noting that illuminating UTBS with photons instead of electrons opens up the space of sample preparation for PEEM imaging. It is plausible that different sample preparations may be even more suitable for PEEM and that combinations of existing EM stains or entirely new stains could increase contrast. Also, for each contrast agent candidate, higher throughput could be achieved by identifying the optimal excitation wavelength. Lastly, UV PEEM is thought of as a surface imaging technique, sampling only the top few nanometers [35]. This makes it an interesting match for gas cluster ion milling [36]. Stain, laser wavelength, and milling thickness could be matched to extract the maximum amount of information from the sample in minimum time. In summary, further exploration of PEEM as a connectomic or projectomic mapping technique can only increase its capabilities and open new pathways for providing large-volume circuit mapping.

## 5 Methods

### 5.1 Sample Preparation

Mouse brain sections were prepared for PEEM imaging following the same protocol used for standard SEM preparation using multiple rounds of osmium tetroxide staining [24]. In brief, the mouse was perfused transcardially to preserve the ultrastructure. The mouse brain was surgically removed, post-fixed, and a vibratome-sectioned brain slice (200 μm to 300 μm) was stained with successive rounds of osmium tetroxide, potassium ferrocyanide, uranyl acetate, and lead nitrate, before ethanol dehydration and embedding with EPON resin. Resin-encapsulated brain was then sliced using an ultramicrotome to 40-80 nm-thick UTBS, which were picked up on a Si substrate (with native oxide termination) coated with 50 nm of gold and glow-discharged.

### 5.2 Photoemission electron microscopy

The UTBS on the substrate is illuminated using a broadband mercury arc lamp with a short-pass filter allowing photons with energies greater than 4.43 eV (280 nm) to focus on the sample, shown schematically in Figure 1. The resulting photoelectrons are imaged with a photoemission electron microscope manufactured by Focus GmbH (Figure 2) and SPECS GmbH (Figure 3 and 4, aberration-corrected microscope). The arc lamp spectrum for the Focus GmbH setup is shown in [37]. The typical exposure time for one image is hundreds of seconds. High-resolution images are obtained by integrating the same region of interest (ROI) to improve signal-to-noise. To estimate how many photons would be required to elicit an electron, we used a pulsed laser with a wavelength of 244 nm and known flux, and acquired a PEEM image of a 60 nm brain slice deposited on gold. We estimated the amount of electrons collected by measuring the SNR ratio and calculating from that shot noise measurement back to the number of electrons collected.

### 5.3 Image processing

2D image montaging was performed in FIJI (ImageJ) using the Grid/Collection stitching plugin. PEEM image stacks were 3D-aligned using AlignTK (https://mmbios.pitt.edu/aligntk-home). To achieve the necessary contrast of UTBS in PEEM, multiple images of the same ROI were captured which introduced drift to the set of images, presumably from instabilities in the stage. Therefore, two people (G.W. and K.M.B.) independently visually inspected the series of images to estimate the pixel drift which was corrected by shifting the images by the estimated pixel drift. Lastly, a maximum intensity projection was generated from the drift-corrected series to produce the final PEEM images. Manual 3D annotation was done using VAST [38].

### 5.4 Space charging simulations

Electrons were simulated to be emitted from a 40 μm × 40 μm substrate with a spatial distribution according to an existing low-noise EM image (10 nm pixel size) from a UTBS of neuropil. They were given an isotropic energy distribution with a standard deviation of 2 eV. Through simulation we determined that suppression of emission due to space charging was a negligible effect even at the highest sample current densities investigated. After emission, the electrons were subjected to a homogeneous electric field of 15 kV/mm. In each 10 fs time step of the simulation, the Coulomb interaction of each pair of in-flight electrons was calculated and applied to the electrons. Once an electron had a distance greater than 1 mm from the sample, it was removed from the simulation. For each job, this process was repeated until 5000 electrons had been removed that were not affected by simulation ramp-up effects. Then, using their final position and angle, each electron was back-projected to the virtual image plane, located 1 mm behind the sample [39], and the electron location in this virtual image plane was compared to the actual emission location. Throughput was calculated using the Rose contrast criterion [26]. All in all 400,000 jobs were executed.

## 6 Acknowledgments

We thank Janek Rieger, Peter Littlewood, Winfried Denk, Rolf Koenenkamp, S. Murray Sherman, John Maunsell, Mark Schnitzer and David Coutts for helpful discussions and/or comments on the manuscript. We thank Jerzy Sadowski, Oliver Schaff, Chris Nicholson, Lisa van Leeuwen, Marcel Hesselberth, Helder Marchetto, and Pupa Gilbert for supporting this project. We thank Vitali Prakapenka and Stella Chariton for help with the laser experiments. This work was partially supported by the University of Chicago Materials Research Science and Engineering Center, which is funded by the National Science Foundation under award number DMR-2011854 and DMR-1420709. This work made use of the shared facilities at the University of Chicago Materials Research Science and Engineering Center, supported by National Science Foundation under award number DMR-2011854. R.L. acknowledges support from a MRSEC-funded graduate research fellowship (DMR-2011854). S.B.K. acknowledges start-up funding support from the University of Chicago and the Neubauer Family Assistant Professors Program. L.L. acknowledges support from the University of Chicago Neuroscience Early Stage Scientist Training Program 1R25NS117360. Part of this project has received funding from the European Union’s Horizon 2020 research and innovation program under grant agreement No 101017902.

## 7 Data availability

Raw data and code will be made available on Zenodo and GitHub upon publication.

---

**Supplementary Figure 1:**
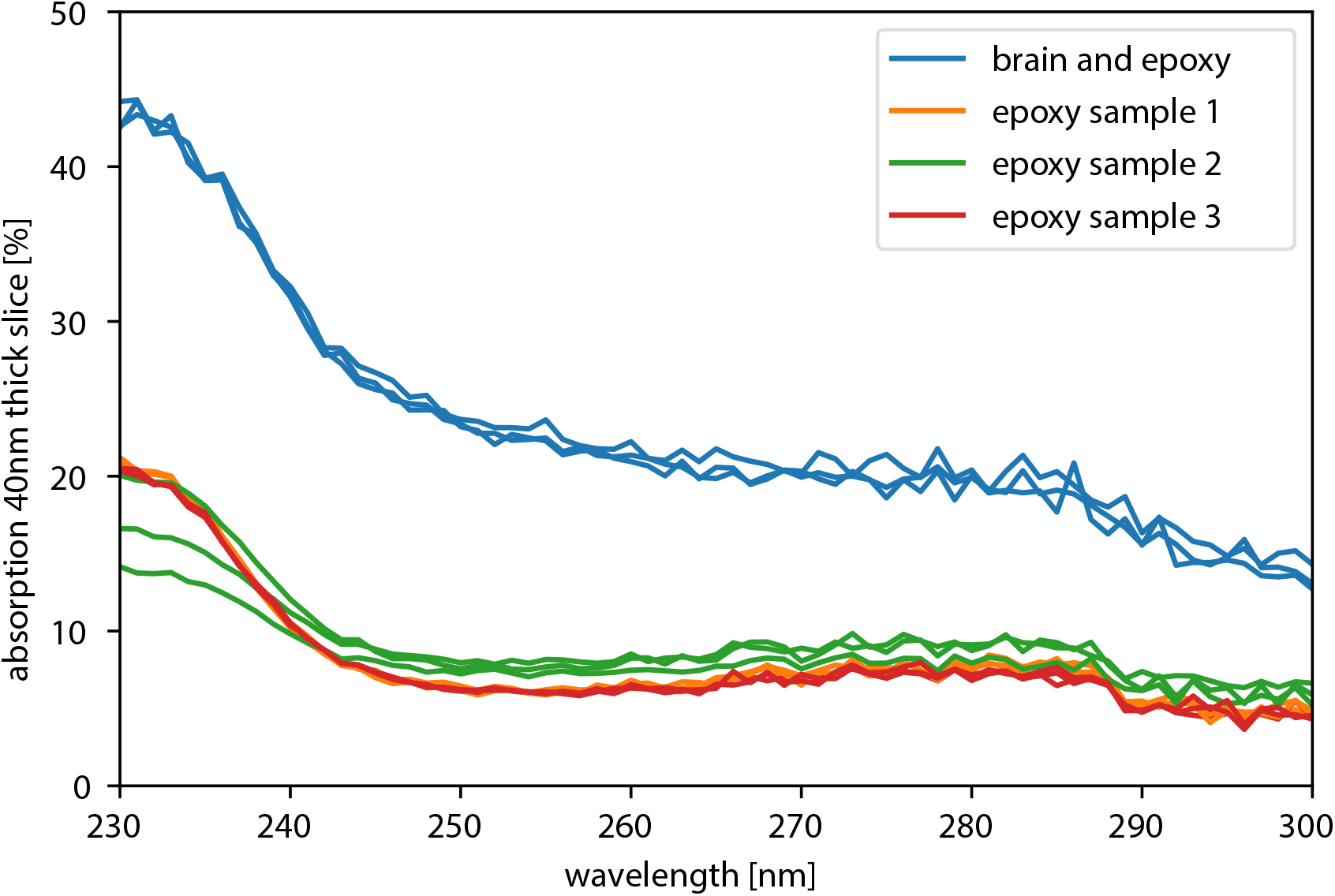
Measurement of the UV absorption of ultrathin slices of epoxy with embedded brain and without. A substantial share of the UV light is absorbed, suggesting an efficient use of laser energy. The upper group of plots shows 3 measurements in 1 sample of epoxy and neuropil, the lower group shows 3 measurements each for 3 samples of pure epoxy.

**Supplementary Figure 2:**
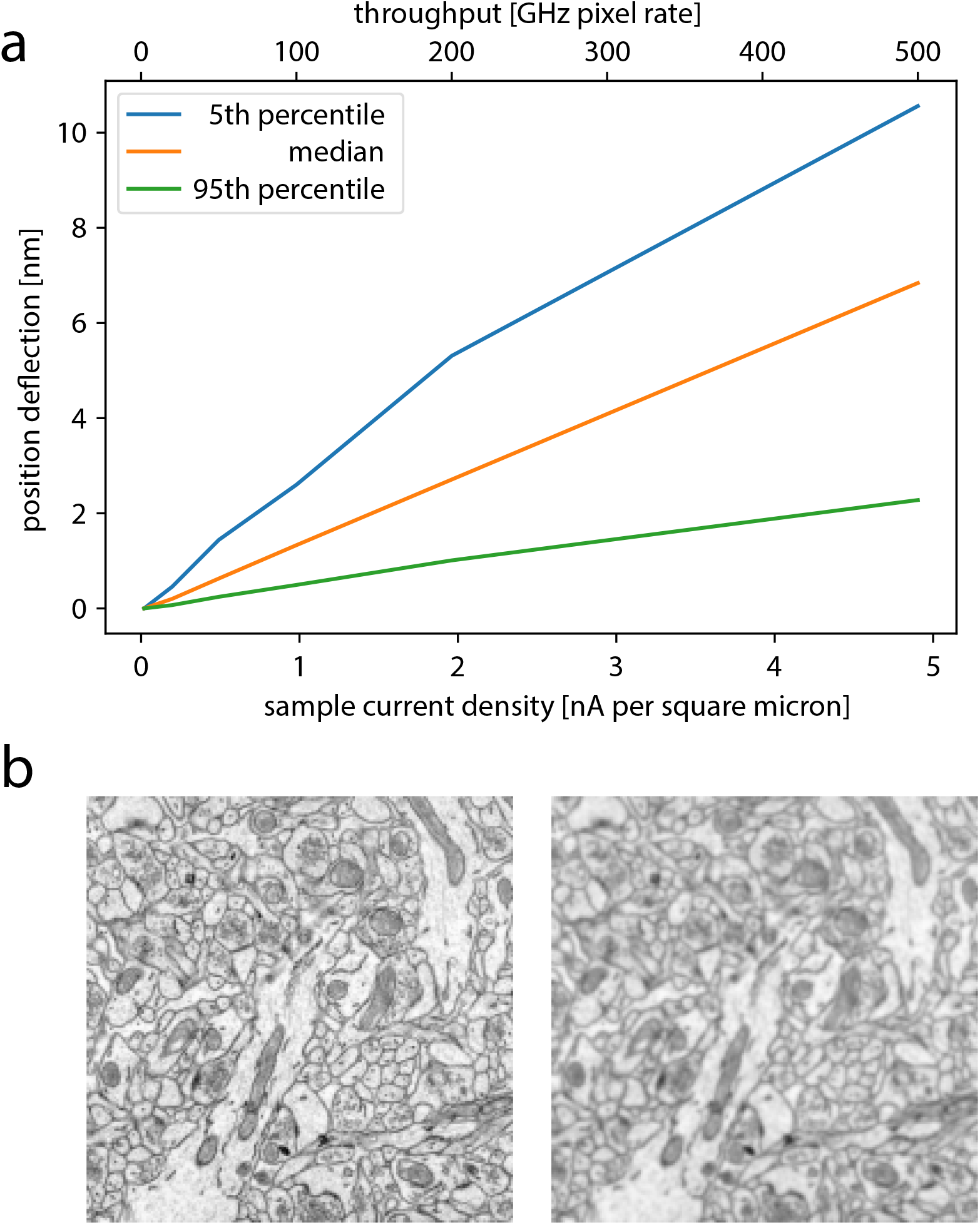
Simulation results estimating space charging effects for high-throughput PEEM imaging for biological materials. A) Electron emission from a UTBS of neuropil and all electron-electron interactions in the space between sample and objective lens were simulated. For more details see Methods section. B) Zoom in (1.8 μm FOV) of example image (high-res TEM) for 0.02 nA μm^−2^ (left, equivalent 2 GHz pixel rate throughput for 40 μm FOV) and 5 nA μm^−2^ (right). Notably, the 2 GHz image is identical to the source image. Also, the results for 200 GHz are plausibly compatible with connectomic imaging. That means that a laser with 1% duty cycle would be suitable for 2 GHz imaging.

**Supplementary Figure 3:**
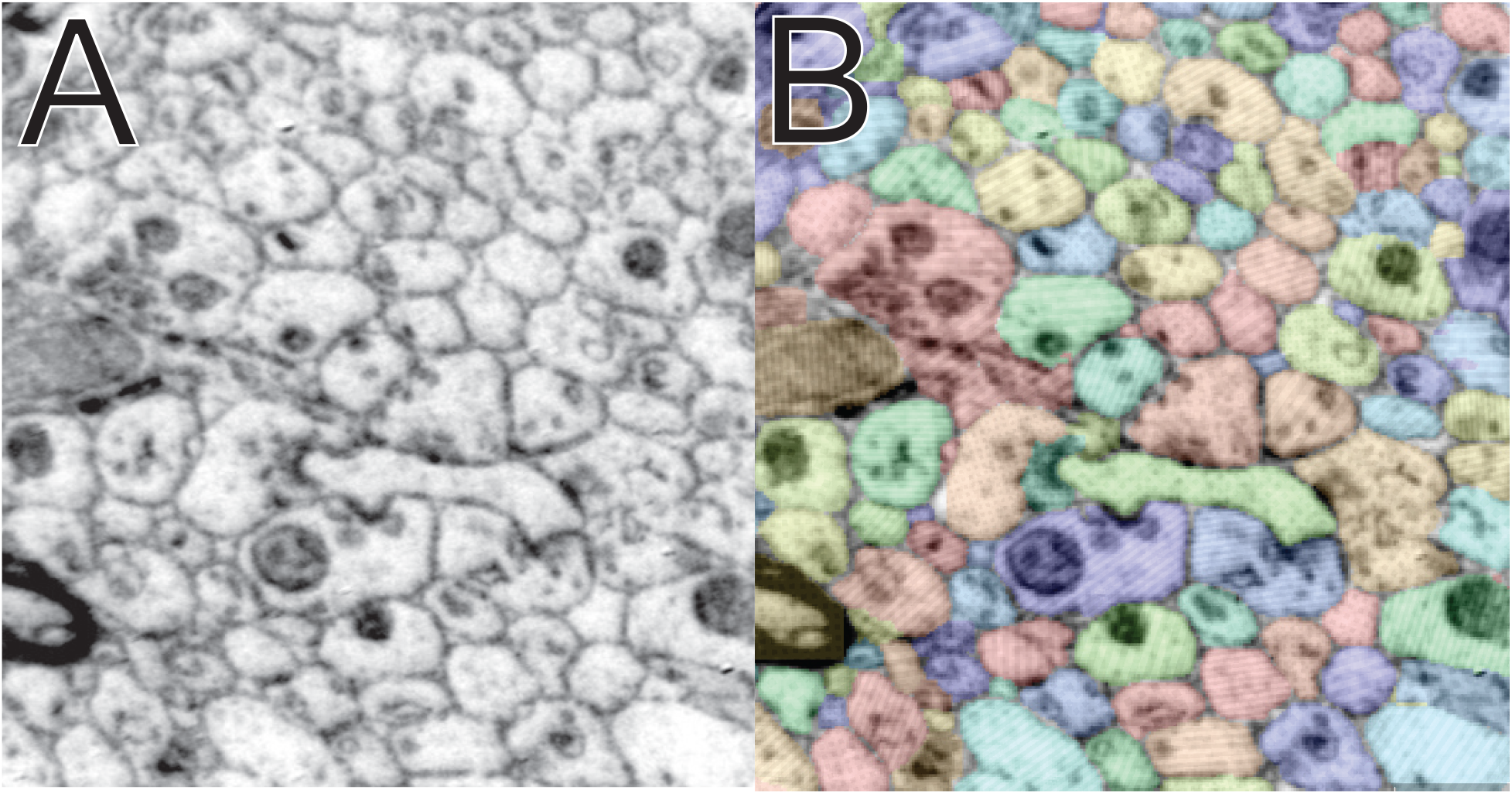
Segmentation of PEEM data: We used the SegmentAnything [30] mode of webKnossos [40] to segment data from Figure 4. With minimal training, SegmentAnything could identify cell boundaries of nearly all cells.

**Supplementary Table 1:**
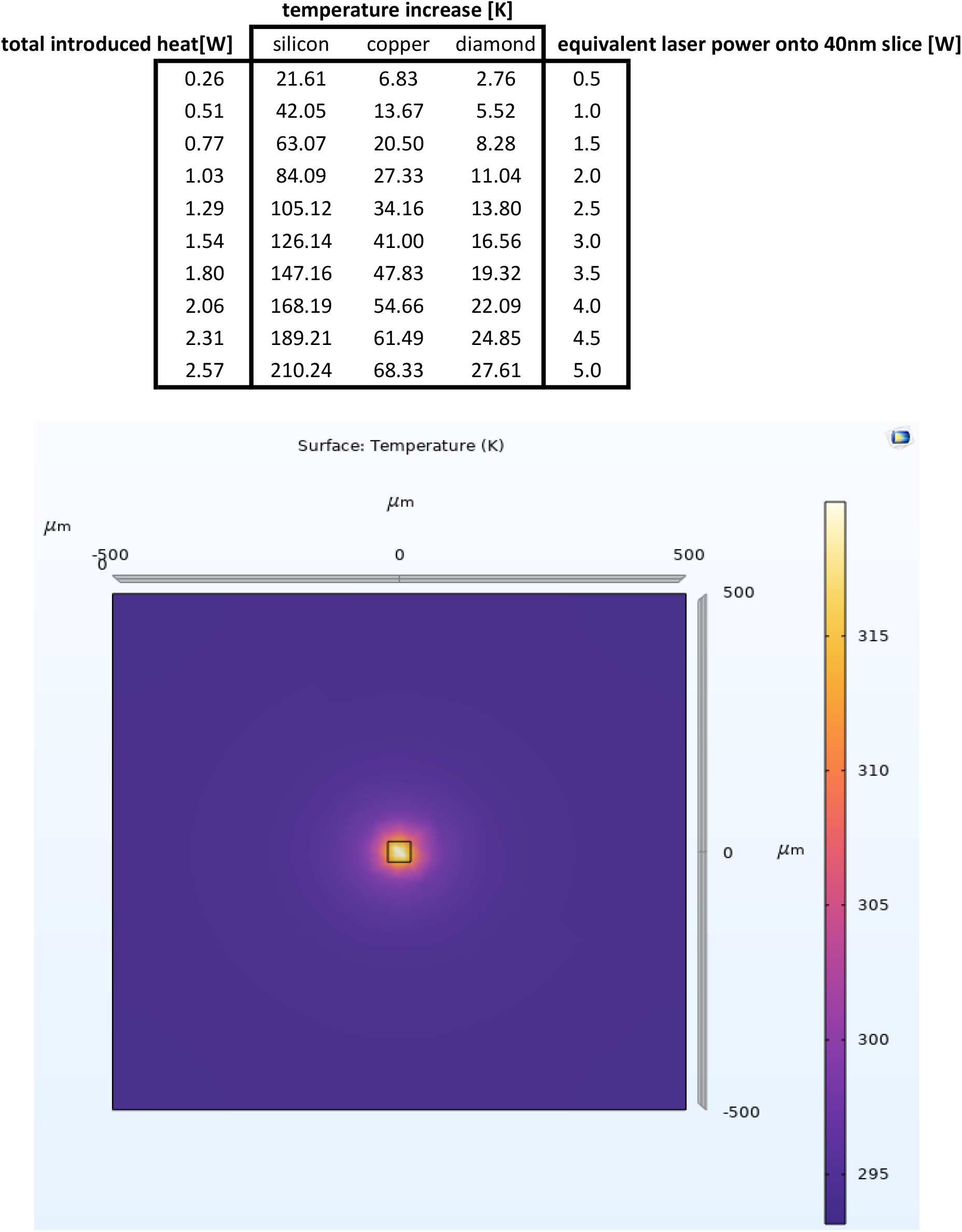
COMSOL simulations of laser heating of UTBS. The heating of brain slices was simulated for a range of laser power settings and for three substrate materials (silicon, copper, and diamond). With increasing thermal conductivity, the substrate is heated less for a given laser power setting.

