## Supplement 1 for "Photoemission electron microscopy for connectomics"

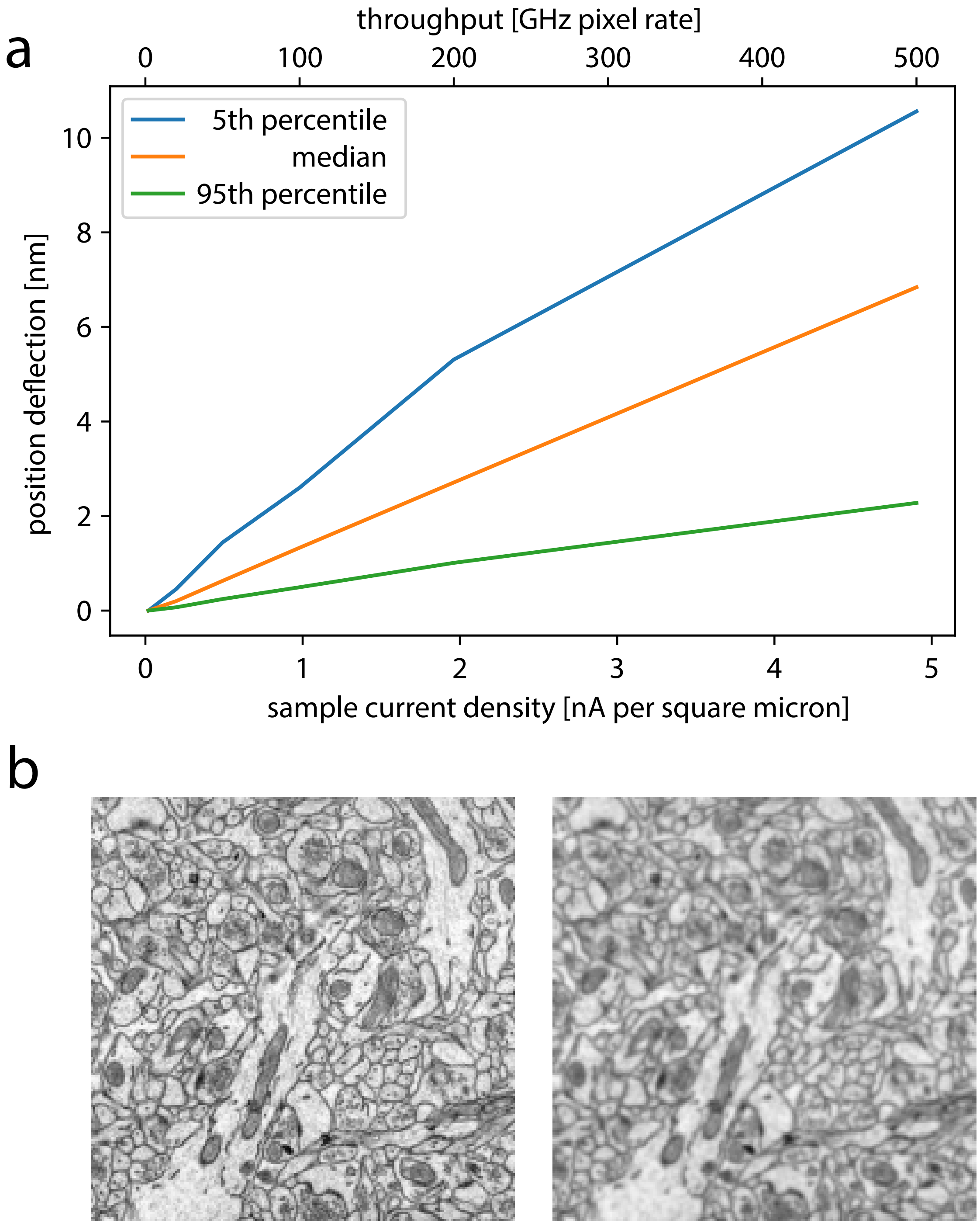


**Supplementary Figure 1: Simulation results estimating space charging effects for high-throughput PEEM imaging for biological materials**. A) Electron emission from a UTBS of neuropil and all electron-electron interactions in the space between sample and objective lens were simulated. For more details see Methods section. B) Zoom in (1.8 µm FOV) of example image for 0.02 nA µm$^{-2}$ (left, equivalent 2 GHz pixel rate throughput for 40 µm FOV) and 5 nA µm$^{-2}$ (right). Notably, the 2 GHz image is identical to the source image.
