## Supplement 2 for "Photoemission electron microscopy for connectomics"

| | temperature increase [K] | | | |
|---|---|---|---|---|
| **total introduced heat[W]** | silicon | copper | diamond | **equivalent laser power onto 40nm slice [W]** |
| 0.26 | 21.61 | 6.83 | 2.76 | 0.5 |
| 0.51 | 42.05 | 13.67 | 5.52 | 1.0 |
| 0.77 | 63.07 | 20.50 | 8.28 | 1.5 |
| 1.03 | 84.09 | 27.33 | 11.04 | 2.0 |
| 1.29 | 105.12 | 34.16 | 13.80 | 2.5 |
| 1.54 | 126.14 | 41.00 | 16.56 | 3.0 |
| 1.80 | 147.16 | 47.83 | 19.32 | 3.5 |
| 2.06 | 168.19 | 54.66 | 22.09 | 4.0 |
| 2.31 | 189.21 | 61.49 | 24.85 | 4.5 |
| 2.57 | 210.24 | 68.33 | 27.61 | 5.0 |

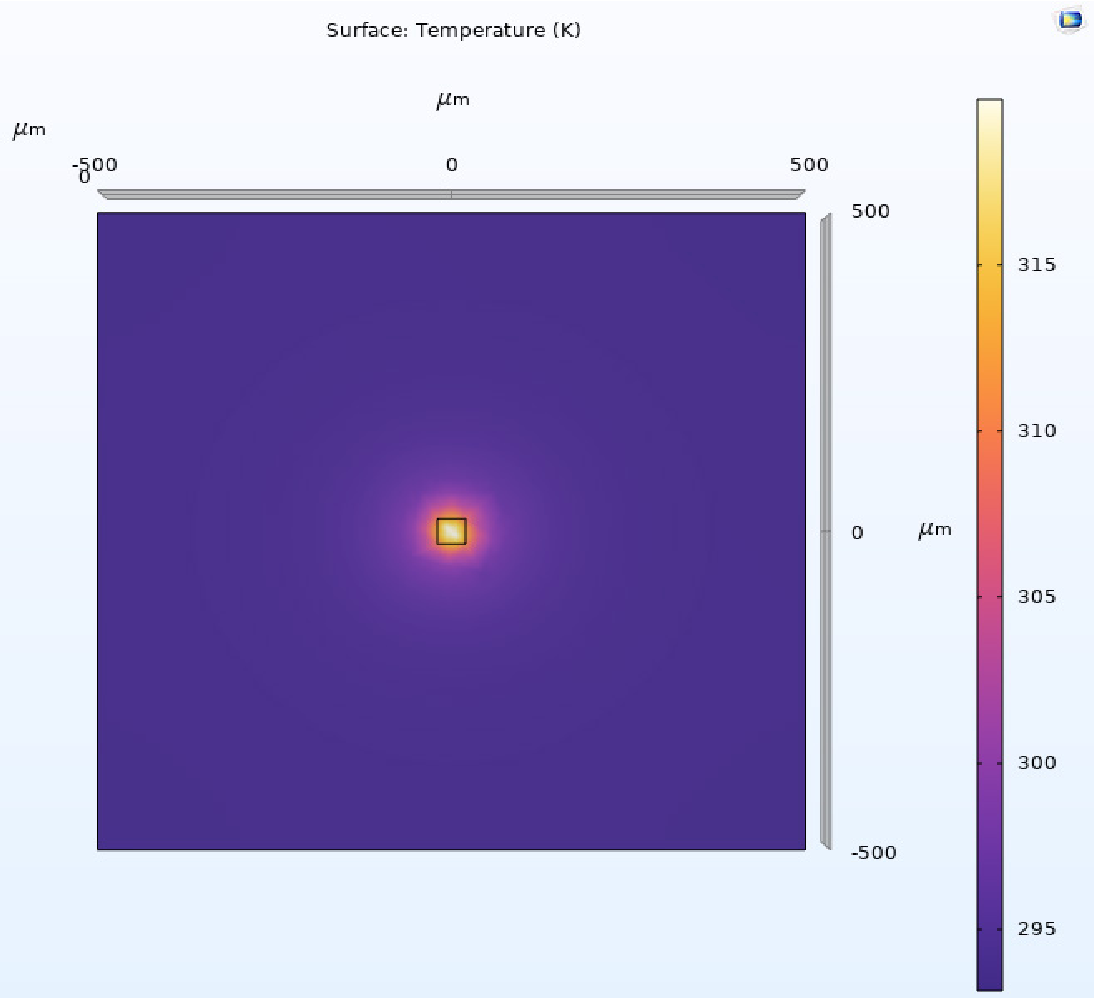


**Supplementary Table 1: COMSOL simulations of laser heating of UTBS.** The heating of brain slices was simulated for a range of laser power settings and for three substrate materials (silicon, copper, and diamond). With increasing thermal conductivity, the substrate is heated less for a given laser power setting.
